# Trynity controls epidermal barrier function and respiratory tube maturation in *Drosophila* by modulating apical extracellular matrix nano-patterning

**DOI:** 10.1101/483768

**Authors:** Yuki Itakura, Sachi Inagaki, Housei Wada, Shigeo Hayashi

## Abstract

The outer surface of insects is covered by the cuticle, which is derived from the apical extracellular matrix (aECM). The aECM is secreted by epidermal cells during embryogenesis. The aECM exhibits large variations in structure, function, and constituent molecules, reflecting the enormous diversity in insect appearances. To investigate the molecular principles of aECM organization and function, here we studied the role of a conserved aECM protein, the ZP domain protein Trynity, in *Drosophila melanogaster*. We first identified *trynity* as an essential gene for epidermal barrier function. *trynity* mutation caused disintegration of the outermost envelope layer of the cuticle, resulting in small-molecule leakage and in growth and molting defects. In addition, the tracheal tubules of *trynity* mutants showed defects in pore-like structures of the cuticle, and the mutant tracheal cells failed to absorb luminal proteins and liquid. Our findings indicated that *trynity* plays essential roles in organizing nano-level structures in the envelope layer of the cuticle that both restrict molecular trafficking through the epidermis and promote the massive absorption pulse in the trachea.

**Summary Statement:** The zona pellucida domain protein Trynity controls the structural organization and function of the apical extracellular matrix in the epidermis and trachea of *Drosophila*.

## Introduction

The skin is one of the largest organs of the body, with various functions including body temperature regulation [1] and sensory-information detection [2], along with its role as a barrier protecting the internal organs from the external environment. The barrier function is provided by the epidermis, which consists of multiple layers: the epidermal cell, stratum corneum, and sebum layers. The outer two layers are non-cellular structures that prevent the entry of external agents such as microbes, viruses, and chemicals, and maintain moisture in the skin [3-5]. Pathogenic conditions that disorganize those layers allow the invasion of external agents through the skin and cause a loss of moisture, resulting in dry skin, asteatosis, and atopic dermatitis [6]. In insects, the body is covered with cuticle layers: the envelope, epicuticle, and procuticle [7]. The cuticle is produced by epidermal cells and provides the barrier function. In both the skin epidermis and cuticle systems, the outermost layers are rich in lipid. The inner layers of the vertebrate epidermis are rich in keratin-family proteins, while the insect epicuticle and procuticle are rich in proteins and polysaccharide chitin, respectively. Thus, these systems share functional and structural similarities.

ZP domain proteins were first identified as structural components of the zona pellucida, the mammalian egg coat [8]. ZP domains are typically ~260 residues long, divided into N-terminal and C-terminal halves with conserved cysteine residues [9], and act as protein-oligomerization modules in the formation of filaments and matrices [10]. ZP domain proteins are reported to function in mammalian-egg fertilization, body-shape regulation, and mammalian auditory-organ formation [reviewed in [11] and [9]].

Twenty ZP domain proteins have been identified in the genome of *Drosophila melanogaster* [12,13], among which *dumpy* (*dpy*) and *piopio* (*pio*) play critical roles in the morphological development of the trachea, wing, and notum [14-16]. In these processes, the ZP domain proteins serve as an anchoring structure that stabilizes the apical plasma membrane of the epidermal tissues undergoing morphogenetic movement [15-18]. ZP proteins also function to shape the denticles, actin-rich apical protrusions in the larval epidermis, by differentially distributing to specific subcellular locations [13]. However, the ZP protein functions are still incompletely understood, in part because many of the available mutant alleles are caused by transposon insertions or large chromosomal deletions. Thus, as a first step toward comprehensively understanding the ZP domain proteins, we performed a systematic mutagenesis of ZP domain proteins in *Drosophila* using the CRISPR/Cas9 genome editing technique (Itakura et al., unpublished). Here we focus on the ZP protein encoded by *trynity* (*tyn*), which was previously implicated in denticle morphogenesis [13]. While this work was in its final stage, it was reported that *tyn* mutants show abnormal feeding behavior and incomplete respiratory-system maturation [19]. The authors suggested that these phenotypes could be explained, in part, by defects of the valve structure of the posterior spiracle. However, previous studies of this gene must be interpreted with caution, because the *tyn* mutant allele were an intronic P- element insertion [*tyn*^*PG38*^in [20] and [19]] or imprecise excisions of the same P element [*tyn*^*ex35*^in [13]]. Thus, it was possible that the phenotypes described in these studies did not represent the loss of Tyn function. Here we used molecularly defined *tyn* null mutations, and demonstrated new roles of *tyn* in the epidermal barrier function and maturation of tracheae that would explain the abnormal feeding behavior, growth retardation, and larval lethality of these mutants. We demonstrated by transmission electron microscopic analyses that *tyn* is required for the formation of the outermost envelope layer of the epidermal cuticle and of pore-like structures in the tracheal cuticle. These results revealed novel functions of *tyn* in constructing nano-level apical extracellular matrix (aECM) ultrastructures.

## Materials and Methods

### Fly strains

Flies were maintained at 25 °C For experiments, adult flies were kept in a vial containing yeast paste on an agar plate overnight. To test viability, 50 L1 larvae on the plate were transferred to a new vial with food, and the number of pupae was counted 7 days later. This process was repeated twice for each genotype. To use embryos or newly hatched larvae for experiments, eggs on the plate were collected, dechorionated in bleach, washed, and kept on a plate with filter paper soaked in water to prevent drying. The dechorionated embryos and hatched larvae at later stages were used. The strains were: *y*^*2*^ *, cho*^*2*^ *, v*^*1*^ *; attP40{nos-Cas9}/ CyO* (NIG: CAS-0001), *y*^*1*^ *,w*^*67c23*^*/P{w+mC=Act- GFP},Dp(1;y)y+* (DGRC:109661), *y*^*1*^ *, w*, baz*^*4*^ *,P{w[+mW.hs]=FRT(w*^*hs*^*)}9- 2/FM7c, P{Dfd-GMR-nvYFP}1, sn*^*+*^(BDSC:23229), w*; P{w[+mC]=His2Av- mRFP1}II.2 (BDSC:23651), *y, w, baz*^*4*^ *,FRT/FM7c, Dfd-GMR-YFP; btl- Gal4,UAS-Serp-CBD-GFP, UAS-p120-tagRFP/CyO, Dfd-YFP* [21,22] and Oregon R (laboratory Stock).

### CRISPR/Cas9 mutant generation

To obtain *tyn* frameshift mutants, we designed a gRNA using a web resource, CRISPR Optimal Target Finder (http://tools.flycrispr.molbio.wisc.edu/targetFinder/) [23] and picked a sequence with a “GG” followed by “NGG” protospacer adjacent motif (PAM) sequence [24] and no off-target matches. The 20-bp target sequence and complementary oligonucleotides with 4-bp overhangs on each end, 5’- CTTCGCGCTCATGGTTCAAATAGG-3’ and 5’- AAACCCTATTTGAACCATGAGCGC-3’, were annealed and cloned into a *Bbsl*- digested gRNA expression vector pBFv-U6.2. This vector was injected into embryos with a genotype of *y*^*2*^ *, cho*^*2*^ *, v*^*1*^ *; attP40{nos-Cas9}/ CyO* using a standard microinjection procedure. Each of 10 founder males was crossed with *y*^*1*^ *, w*, baz*^*4*^ *, P{w[+mW.hs]=FRT(w*^*hs*^*)}9-2/FM7c, P{Dfd-GMR-nvYFP}1, sn*^*+*^, and ~100 independent candidate mutant lines were established. Eleven lines were subjected to a heteroduplex mobility assay (HMA) in 15% acrylamide gel [25], and three of these lines showed a mobility shift. Sequencing revealed two lines with a deletion and consequent stop codon (*tyn*^*1*^and *tyn*^*2*^) and 1 line with a 5-bp replacement. The primers used for HMA were: 5’- GGATCATAAGACCCTGCCCG-3’, 5’-TGGTGATCCAGCTCCCAAAC-3’, and for sequencing: 5’-CCGAAGAGAAGGTTGCCCAA-3’, 5’- TGCGGGTTAAGTTGGTCAGG-3’. We also picked one established line with no mutation as a control.

### Histology

For cuticle preparations, larvae were placed on a glass slide with Hoyer’s medium and lactic acid (Wako) 1:1 and cover glass, and incubated at 60-65 °C overnight. Images were obtained with an Axioplan2 (Zeiss) and DP74 camera (Olympus). Eosin staining was performed with a modified protocol of Zuber et al. [26]. Intact L1 larvae were incubated in 0.5% Eosin Y staining solution (Sigma-Aldrich) for 1 hour at 40 °C and washed with water. After heat fixation at 70 °C for ~1 min [27], the larvae were observed with a VHX-6000 digital microscope (Keyence). For the DAPI penetration test, DAPI (Sigma) in water (1 μg/ml) was applied to intact larvae in 50 μl of PBS [10X Phosphate-Buffered Saline (Nacalai Tesque) diluted with water] and washed out 5 min later. To image the DAPI-immersed larvae or larvae expressing Serp-CBD-GFP and p120-tagRFP for the protein clearance test, the samples were mounted in glycerol, fixed by heating, and observed with a Fluoview FV1000 confocal microscope (Olympus).

### Physiological tests

To examine the effects of low osmotic pressure, dechorionated embryos were placed in water overnight. We picked 10 larvae for each genotype and calculated the survival rate and relative body lengths of the surviving larvae normalized to the median obtained for Oregon R. Since high osmotic pressure caused rapid changes, 10 hatched larvae were collected in 5 μl PBS, and 20 μl 10X PBS was added. Thirty minutes later, images were captured and survival rates were calculated.

### In situ hybridization

Because the probe used previously [12] (1082-bases long) showed relatively high background signals in our experiments, we designed a new probe that was 356-bases long. This new probe detected the *tyn* mRNA expression in a pattern similar to that reported previously, but the lower background enabled us to detect signals in the tracheal system more clearly. The digoxygenin-labeled RNA probe was synthesized with T7 RNA polymerase and a template amplified with the primers 5’-GGCGGCCTTTAGTTTTGTGG-3’ and 5’- TAATACGACTCACTATAGGGGTCCAGAGCTGCGTCTATCC-3’ (with T7 promotor) followed by gel filtration. Whole-mount *in situ* hybridization was performed as described previously [28]. Embryos were mounted in 80% glycerol, observed using a BX53 upright microscope with DIC optics, and photographed with a DP74 digital camera (Olympus).

### Time-lapse Imaging

To observe feeding behavior, 10 larvae were placed in each well of a 96-well plate with agar, and yeast paste was applied to the center of each well. Time-lapse imaging was then performed at 15-sec intervals, for a total of 2 hours. Immobile larvae were excluded. The percentages of larvae inside, peripheral to, and outside the paste were plotted against time after the paste was applied. For live imaging of the gas-filling process, stage-17 embryos with visible malpighian tubules and slightly pigmented mouth cuticle, most of which should enter the stage of tracheal liquid-gas transition within an hour, were picked and placed on a plate with heptane glue (heptane with sticky tape) and covered with water. Images were obtained every 30 sec. for 6 hours. For both imaging experiments, a Digital Microscope VHX-6000 (Keyence) was used.

### Transmission electron microscopy

The TEM protocol was modified from a previously described procedure [29]. Dechorionated embryos at the same stage as those used for live imaging of the liquid-gas transition were placed on sticky tape attached to a glass slide, poked out from the vitelline membrane in fixation solution (2% paraformaldehyde, 2% glutaraldehyde, 1xPBS, 1 mM CaCl_2_, 1.5% DMSO, pH=7.4), transferred to fresh fixation solution, and further incubated for 2 hours in total. The embryos were washed with PBS, embedded in 3% agarose, and cut out as agarose cubes containing an embryo to make the following processes easier. After the subsequent fixation (2% OsO4, 1x PBS, 0.2 M sucrose) on ice for 2 hours and dehydration with a graded series of ethanol in water (30% and 50% on ice, 70%, 80%, 90%, 95%, 99.5%, 100% 3 times, at room temperature, 15 min for each), the embryos were embedded in resin (propylene oxide for 20 min twice, propylene oxide-resin 1:1 for 1 hour, 1:3 overnight, 100% resin for 3 hours twice, and overnight under vacuum) and incubated at 60 °C for polymerization. Ultrathin 80-nm-thick sections were obtained and stained with heavy metal. Images were captured with a JEM-1400 (JEOL). Three embryos each for *y*^*2*^ *, cho*^*2*^ *, v*^*1*^and *tyn*^*1*^ *, y*^*2*^ *, cho*^*2*^ *, v*^*1*^were examined.

## Results

### *tyn* mutants exhibit growth and behavioral defects

Trynity has three PAN/Apple domains followed by a ZP domain and a transmembrane domain (Fig. 1A). We mutagenized *tyn* using the CRISPR/Cas9 technique [30,31] by targeting the guide RNA to the sequence encoding the third PAN/Apple domain (Fig. 1A, arrowhead). Two independent mutations with small deletions that resulted in a frameshift and a stop codon were isolated (*tyn*^*1*^and *tyn*^*2*^, Fig.1A). The two mutations were lethal as hemizygous, homozygous, and trans-heterozygous conditions, and this lethality was rescued by a chromosomal duplication carrying the *tyn* locus (Materials and Methods). In the following experiments, we used *tyn* stocks balanced with *FM7-dfd-YFP,* and progenies lacking the dfd-YFP signal (*tyn* homozygotes and hemizygotes) were used as mutants, while animals with the dfd-YFP signal were used as sibling controls. We also used the Oregon R strain, and another control line, *y*^*2*^ *,cho*^*2*^ *,v*^*1*^, which carries a normal *tyn* sequence obtained from the same progeny that yielded *tyn*^*1*^and *tyn*^*2*^, as controls.

**Figure 1:**
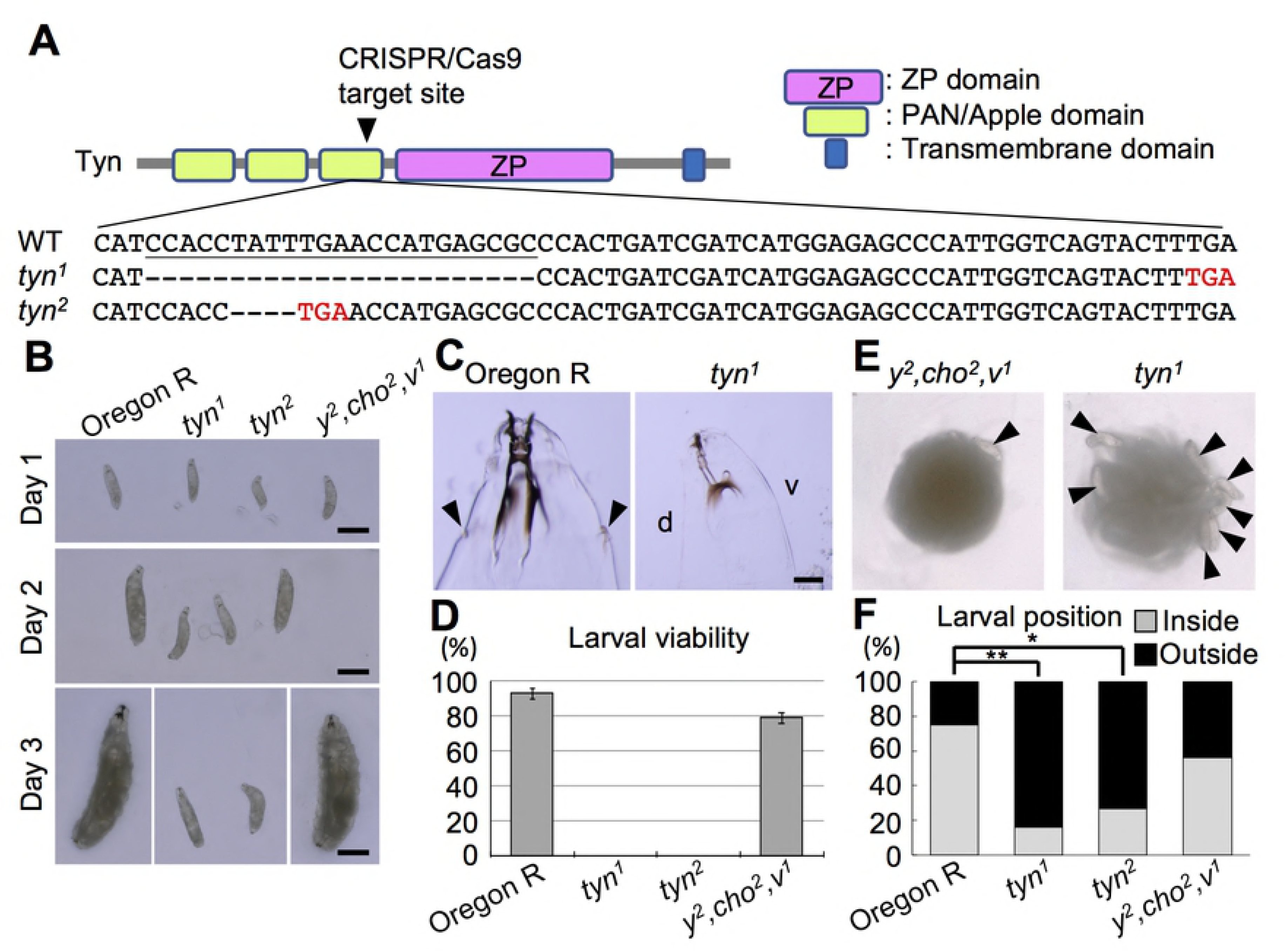
*tyn* mutants and their gross phenotypes. (A) Tyn protein structure and *tyn* DNA sequence of the region including the CRISPR/Cas9 target site. Tyn has 3 PAN/Apple domains, a ZP domain, and a transmembrane domain. The target site is indicated by an arrowhead in the diagram, and the sequence is underlined. *tyn*^*1*^and *tyn*^*2*^contained 23- and 4-base deletions respectively, which resulted in early stop codons (red letters). (B) Larval body size for each genotype. Day 1 means ~24 hours after hatching. On days 2 and 3, the *tyn* mutants remained small, while control larvae (Oregon R and *y*^*2*^ *, cho*^*2*^ *, v*^*1*^) grew larger. (C) On day 3, the *tyn*^*1*^mutant larvae lacked anterior spiracles, while those of Oregon R were obvious (arrowheads). (D) Larval viability of each genotype. n=100 for each. (E) Photographs of larvae with yeast paste. Arrowheads indicate larvae that stayed at the periphery of the yeast paste without entering it. In the left panel, there were 5 larvae within the food. The quantified data are shown in (F). n=15-20 for each. **P<0.01, *P<0.05 by Fisher’s exact test with Benjamini & Hochberg correction.

The *tyn* mutant first-instar larvae hatched with a normal appearance, but their growth was retarded (Fig.1B). On day 3, the mutant larvae lacked anterior spiracles and their head skeletons were underdeveloped, suggesting molting failure (Fig. 1C). All of the mutants died before pupation (Fig.1D).

Both control and *tyn* mutant L1 larvae placed on agar plates approached distantly placed yeast food (Supplementary Fig. 1) and ingested it, as shown by gut labeling by food containing blue dye (data not shown). However, the *tyn* mutants tended to stay at the periphery of the food, while control larvae entered it (Fig. 1E, F and Supplementary Fig. 1). These defects in growth and feeding behavior, and additional defects in posterior spiracle valve formation (Supplementary Fig. 2) were similar to phenotypes reported for *tyn*^*PG38*^[19].

**Figure 2:**
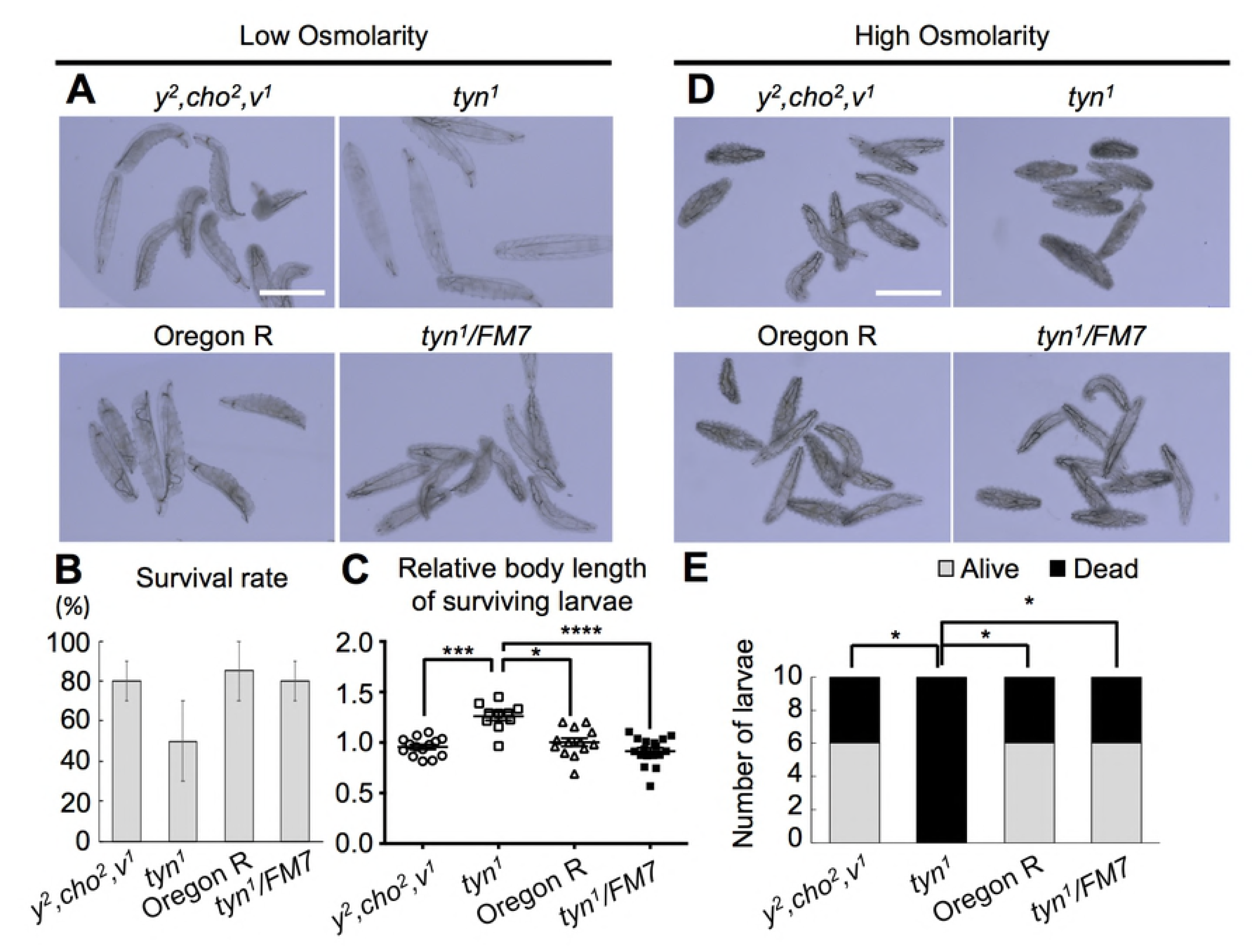
Defective barrier function against osmolarity in *tyn* mutants. First-instar *tyn*^*1*^ *, y*^*2*^ *, cho*^*2*^ *, v*^*1*^larvae showed obvious swelling in water (A-C) and shrinkage in hyper-osmotic solution (D, E) compared with control groups, *y*^*2*^ *, cho*^*2*^ *, v*^*1*^, Oregon R, and *tyn*^*1*^ *, y*^*2*^ *, cho*^*2*^ *, v*^*1*^ */FM7*. (A) Larvae kept in water overnight. (B) The survival rate was calculated by dividing the number of moving larvae by that of total larvae (10 larvae for each of 2 trials, average values ± S.E.M are shown.). There was no significant difference (P=0.29 by Kruskal Wallis test). (C) Median body length normalized to that of Oregon R. ****P<0.0001, ***P<0.001, *P<0.05 by Dunn’s multiple comparisons test after a Kruskal-Wallis test (P<0.0001). (D) Larvae kept in high-salt solution for 30 min. (E) All 10 of the *tyn*^*1*^ *, y*^*2*^ *, cho*^*2*^ *, v*^*1*^larvae died, while 6 of the 10 larvae survived in each of the three control groups. *P<0.05 by Fisher’s exact test with Benjamini & Hochberg correction. Scale bar: 500 μm.

### *tyn* is essential for the epidermal barrier function

We found that the *tyn*^*1*^mutants were sensitive to low osmolality. When first- instar larvae were kept in water overnight, many of the mutant larvae showed dramatic swelling of the body, whereas control larvae did not (Fig. 2A). Although the viability of the mutants was slightly lower in this condition (Fig. 2B), the body length of the surviving *tyn*^*1*^mutant larvae was 1.3-times longer than that of controls (Fig. 2C). On the other hand, incubation under high osmolarity (1 M NaCl) for 30 min. caused prominent shrinkage of the mutant larval body, and a significantly higher death rate than in the three control groups (Fig. 2D and E, p<0.05). These observations indicated that the *tyn*^*1*^mutant larvae were sensitive to changes in the external salt concentration, suggesting that their epidermis allowed the passage of salt ions and water.

We next tested the permeability of the epidermis to larger molecules. Incubating L1 larvae in Eosin Y (molecular weight 647.89) at 40°C for 60 min. resulted in intense staining of the internal organs of *tyn*^*1*^and *tyn*^*2*^mutants but not of control larvae (Fig. 3A, *tyn*^*2*^: data not shown). In addition, incubation in DAPI (molecular weight 350.25) caused the labeling of epidermal cells and tracheal cells of *tyn*^*1*^mutants, but not of control larvae (Fig. 3B). Wang et. al. demonstrated that Eosin Y penetrated the tracheal lumen through permanently open spiracles in *tyn*^*PG38*^larvae, but they did not observe labeling of the tracheal cells, epidermis, or internal tissues [19]. These differences in the *tyn*^*1*^and *tyn*^*PG38*^phenotypes may reflect the different nature of the mutations. Our findings collectively indicated that in the *tyn-*null larvae, the epidermal barrier function failed and allowed external salt and small molecules to leak into the inner tissues.

**Figure 3:**
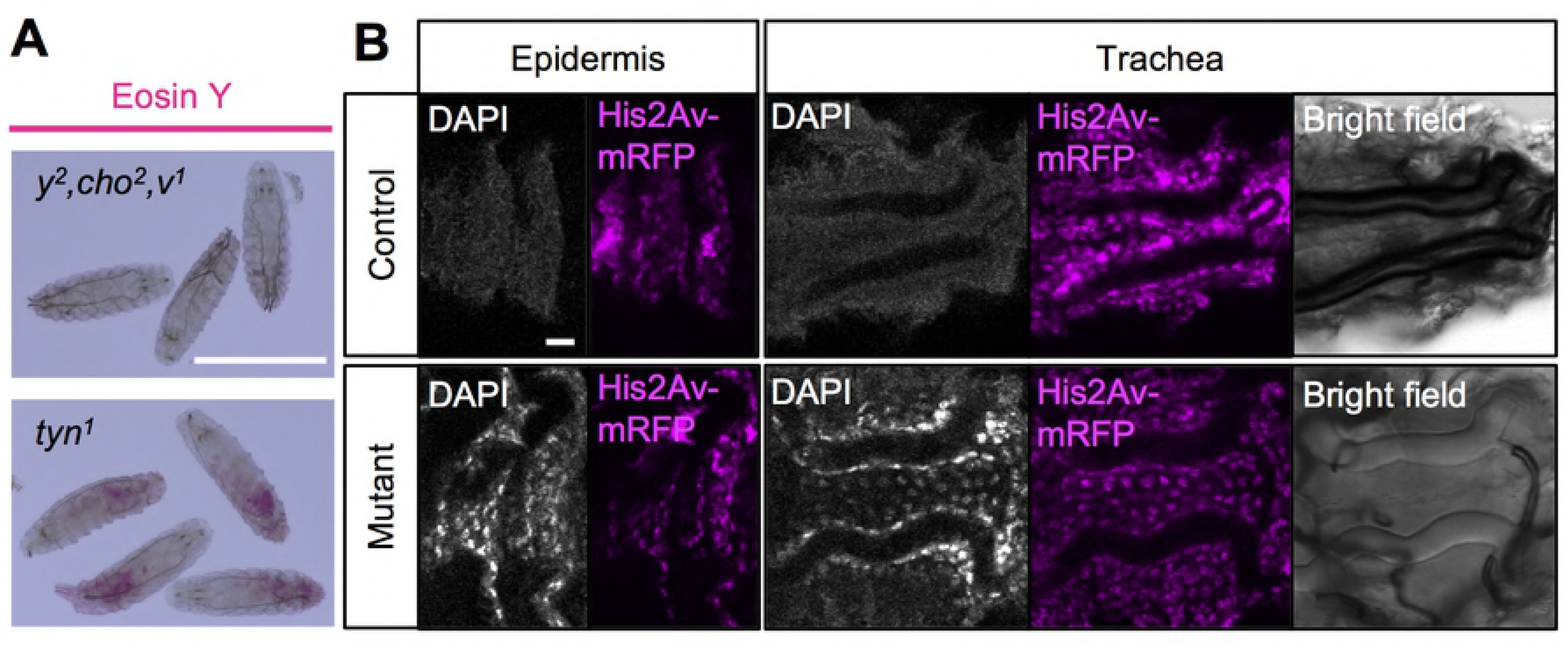
Leakage of molecules through the *tyn* mutant body surface. (A) Eosin Y penetrated the body of *tyn*^*1*^ *, y*^*2*^ *, cho*^*2*^ *, v*^*1*^larvae but not of *y*^*2*^ *, cho*^*2*^ *, v*^*1*^. Scale bar: 500 μm. (B) DAPI staining was observed in neither the epidermis nor trachea of the control larvae (+/+; Ubi-GAP-43GFP, His2AV-mRFP), while some of the epidermal and tracheal cells were strongly labeled with DAPI in the mutant larvae (*tyn*^*1*^ *, y*^*2*^ *, cho*^*2*^ *, v/Y;* Ubi-GAP-43GFP, His2AV-mRFP). His2AV- mRFP (magenta) was expressed to visualize nuclei in all the cells. Bright-field images show that the trachea was gas-filled in the control but not in the mutant. Scale bar: 10 μm.

### The outermost layer of the epidermis is disorganized in *tyn* mutants

To determine the cause of the decreased barrier function in *tyn* mutants, we observed the epidermal structures by transmission electron microscopy (TEM). Moussian et. al. described three major layers of the larval cuticle that are distinguished by TEM: the procuticle, epicuticle, and envelope [32] (Fig. 4A, A’). The procuticle, which mainly consists of chitin and protein, borders the apical side of the epidermal cells. The epicuticle is the next outer layer and rich in proteins. The envelope is the outermost of the three layers, and contains lipids, waxes, and proteins. TEM observation of *tyn*^*1*^mutants at the late stage 17 of embryogenesis showed a three-layered cuticular organization indistinguishable from that of controls, except for the outer layer of the envelope, which was rough and disorganized compared to the smooth control envelope (Fig. 4B, B’, arrows indicate defective structures). Specifically, while the envelope layer of control embryos consisted of five alternating electron-dense and lucid sublayers (Fig. 4C), in the *tyn*^*1*^mutant the surfaces of one or two layers were broken into debris (Fig. 4D).

**Figure 4:**
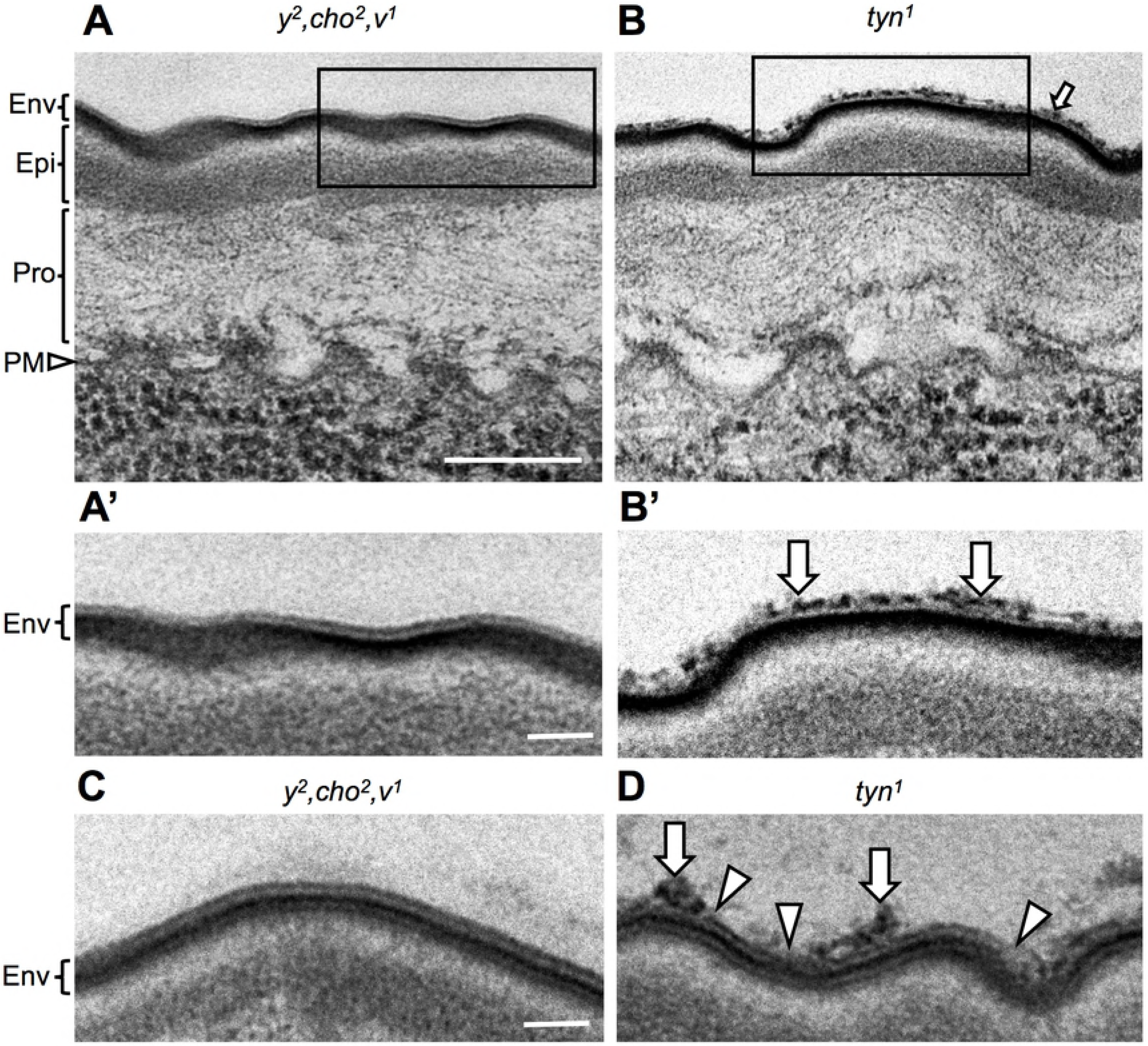
Ultrastructure of the epidermis in *y*^*2*^ *, cho*^*2*^ *, v* and the *tyn*^*1*^mutant. (A) Three cuticle layers formed on the plasma membrane of epidermal cells (PM, arrowhead): the outermost envelope (Env), intermediate epicuticle (Epi), and innermost procuticle (Pro). (B) The surface of the envelope layer of the *tyn*^*1*^mutant was disorganized (arrow). (A’), (B’): Enlarged views of the region indicated by a rectangle in (A) and (B), respectively. In control samples, the five sublayers constituting the envelope were observed clearly, as alternating electron-dense and lucid layers in (C). (D) In the *tyn*^*1*^mutant, one or two surface sublayers were broken in many regions (arrowheads), causing a rough surface with extensive debris (arrows). Scale bar: 200 nm for (A), (B) and 50 nm for (A’), (B)’, (C), (D).

### *tyn* mRNA localizes to the epidermis, trachea, and other organs

We next performed whole-mount embryonic *in situ* hybridization analysis with a newly designed RNA probe for *tyn* (see Materials and Methods). This analysis confirmed the previously reported expression of *tyn* in the mouth parts including the pharynx and esophagus, posterior spiracles (PS), denticle belts, and epidermis (Fig. 5A, C, D, E) [12], as well as in the hindgut (Fig. 5A). In addition, we observed strong *tyn* expression not only in PS but in the whole tracheal system, from stage 14 to stage 17, with a peak at stage 15 (Fig. 5A, B, E, F). The *tyn* mRNA was localized to the apical cortex of the tracheal cells and fibrous structures in the lumen (arrowheads in Fig. 5B’ and F). We speculate that the luminal *tyn* mRNA was co-secreted during a massive secretion pulse of proteins and polysaccharides that is known to occur during tracheal development [33]. The *tyn* expression in the mouth parts and epidermis was strong at stage 16 (Fig. 5C, D). The *tyn* expression in PS and DT lasted until stage 17 (Fig. 5E, F).

**Figure 5:**
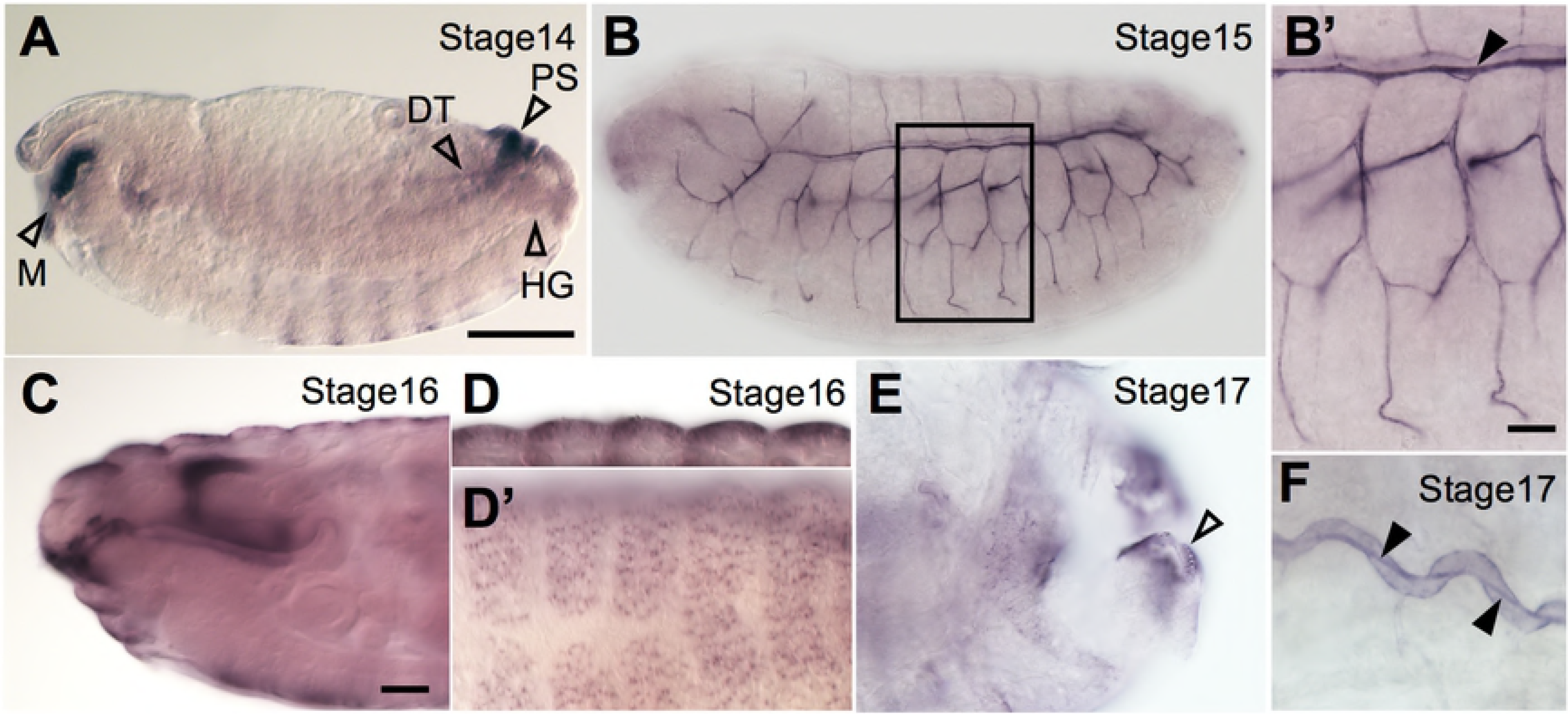
*tyn* mRNA expression at late embryonic stages. (A) At stage 14, *tyn* mRNA was expressed in the posterior spiracles (PS), mouth parts (M), hindgut (HG), and part of the dorsal trunk (DT). (B) The *tyn* mRNA was expressed in the tracheal system most strongly at stage 15. At stage 16, strong expression was observed in the mouth parts (C) and epidermis (D, dorsal view). *tyn* mRNA expression lasted until stage 17 in the PS (E, dorsal view) and DT (F), but not in the epidermis. Scale bar: 100 μm for A, B, 20 μm for B’, C-F. Black arrowheads in B’ and F indicate luminal mRNA.

### *xtyn* is required for protein clearance and gas filling in the tracheal lumen

We next examined the maturation process of the trachea. The *tyn*^*1*^and *tyn*^*2*^mutant larvae lacked fully gas-filled trachea (Fig. 6A), and their posterior spiracles had an abnormal morphology (Supplementary Fig. 2). Based on similar observations in *tyn*^*PG38*^mutants, Wang et al. suggested that the *tyn* mutation caused a loss of posterior spiracle valve structures and unrestricted liquid flow between the tracheal lumen and outside liquid, resulting in a defect in liquid clearance, or gas filling [19]. However, given the strong *tyn* mRNA expression in the whole tracheal system, we sought an alternative explanation for the gas-filling defect. Time-lapse imaging with control embryos revealed the normal time course for gas filling (Fig. 6B i-vi). After gas bubbles initially formed in the central metameres, the gas spread into the dorsal trunk (DT) (Fig. 6B, ii), traversed across dorsal branch 10 (DB10, Fig. 6B, iii), spread into the contralateral DT, and finally filled the other small branches and posterior spiracle (Fig. 6B, iv, white and yellow arrowheads, respectively). In the *tyn* mutants, the gas-filling defect was nearly complete in the posterior spiracle, but was variable in the DT (Fig. 6C). The timing of the first bubble formation in the *tyn* mutants showing full or partial gas filling was delayed by ~3 hours compared to control embryos (Fig. 6D), while the duration of the hatching behavior was normal (Fig. 6E).

To explore the reason for the significantly delayed onset of gas bubble formation, we observed the events prior to tracheal tube maturation. The tracheal branching, fusion, tube elongation, and diameter expansion proceeded normally in the *tyn* mutants, resulting in a DT filled with chitin at stage 16 (data not shown). In normal embryos, a massive absorption of luminal proteins occurs at stage 17, followed by gas-filling or liquid clearance (Fig. 6F;[33]). We found that *tyn* mutants at stage 17 retained a luminal protein marker [GFP-tagged Serpentine Chitin Binding Domain (Serp-CBD-GFP) [21]], suggesting that the endocytosis of luminal proteins failed (Fig. 6G). These observations indicated that the *tyn* mutant trachea had a defect in protein clearance, resulting in delayed and incomplete gas-filling.

**Figure 6:**
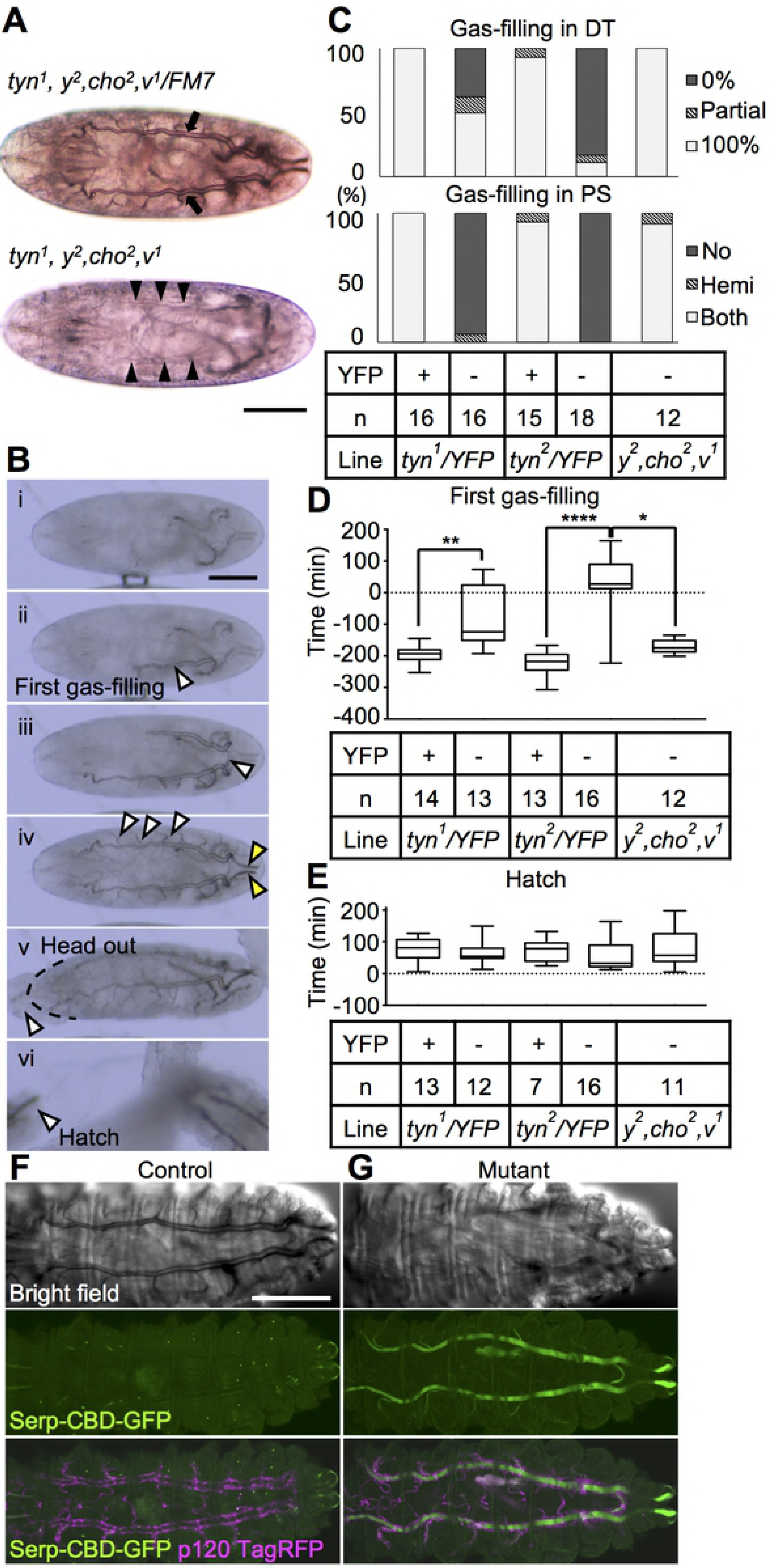
Delayed and partial gas-filling, and protein clearance failure in *tyn* mutants. (A) Tracheal tubules in a *tyn*^*1*^sibling control embryo were filled with gas and bordered by a thick black line (upper panel, arrows). In the *tyn*^*1*^mutant, the trachea had translucent borders and lacked gas (bottom panel, arrowheads). (B) The normal gas-filling process observed in a *tyn*^*1*^sibling control: (i) tracheal tube before liquid-gas transition, (ii) first gas-filling, (iii) extension to the contralateral side through posterior anastomosis (arrowhead), (iv) subsequent propagation to the branches (white arrowheads) and posterior spiracles (yellow arrowheads), (v) extrusion of the head from the vitelline membrane, (vi) hatching. (C) The degree of gas-filling was classified into three levels for each of the two parts, dorsal trunk (DT, upper) and posterior spiracles (PS, bottom), for each of five genotypes: *tyn*^*1*^sibling control, *tyn*^*1*^mutant, *tyn*^*2*^ sibling control, *tyn*^*2*^mutant, and no-mutation control (left to right). The YFP signal indicated the presence of the X-chromosome balancer, FM7 with Dfd-YFP. (D, E) The time when the embryo’s head emerged from the vitelline membrane (B-v) was set as 0, and the timing of the first gas-filling (D) or hatching (E) is shown as a boxplot with S.E.M. for each of the five genotypes. For the first gas-filling time, a Kruskal-Wallis test followed by Dunn’s multiple comparisons test detected significant differences between the three pairs: *tyn*^*1*^ sibling control vs mutant (**P<0.01), *tyn*^*2*^sibling control vs mutant (****P<0.0001), and *tyn*^*2*^mutant vs no-mutation control (*P<0.05). There was no difference in the hatching time. (F, G) First-instar larva expressing Serp-CBD-GFP (green) and p120-TagRFP (magenta) by the btl-gal4 driver with (F) or without (G) a functional *tyn* gene. (F) The control tracheal tube was filled with gas and Serp-CBD-GFP was completely removed from the lumen. (G) The *tyn* mutant lumen was filled with Serp-CBD-GFP. Scale bar: 100 μm.

### Pore-like structures in the tracheal cuticle are defective in *tyn* mutants

Prior to the protein and liquid clearance, cuticle deposition begins at stage 16, and involves the formation of taenidial folds over the apical cell surface of tracheal cells [34]. Since the cuticle layers physically separate the luminal liquid and molecules from the plasma membrane of tracheal cells, how the massive pulse of luminal liquid and molecule absorption occurs is not well understood. To examine this process, we used TEM to observe the ultrastructure of the tracheal cuticle. In control flies, the longitudinal section of the DT exhibited regularly spaced ridges of taenidial folds, which were separated by thin interteanidial cuticle that was enriched with electron-dense materials (Fig. 7A). These materials occasionally appeared as parallel dense lines, indicating a pore connecting the luminal space to the cell surface (Fig. 7A, A’, A”, arrowheads). The inner space was lucent and around 20-nm in diameter. Hereafter we will call this pore structure a “taenidial channel” (TC). In the *tyn*^*1*^mutants, the electron-dense materials of the TC were cloudy, and distinct TCs were not observed (Fig. 7B, B”). In addition, the apical tracheal cell membrane was slightly bulged out at the site of intertaenidial cuticle in the control (Fig. 7A, A’), and this bulging was more prominent in the *tyn*^*1*^mutants (Fig. 7B’, arrow).

## Discussion

In this study, using newly established *tyn-*null mutants, we showed that *tyn* is required for the building of specific substructures in the epidermal and tracheal cuticles. The *tyn* mutations caused defects in epidermal barrier function and in luminal protein clearance of the tracheal tubule, resulting a variety of physiological and behavioral problems. Based on these results, we were able to elucidate some of the physiological roles of cuticular substructures, which are discussed below.

**Figure 7:**
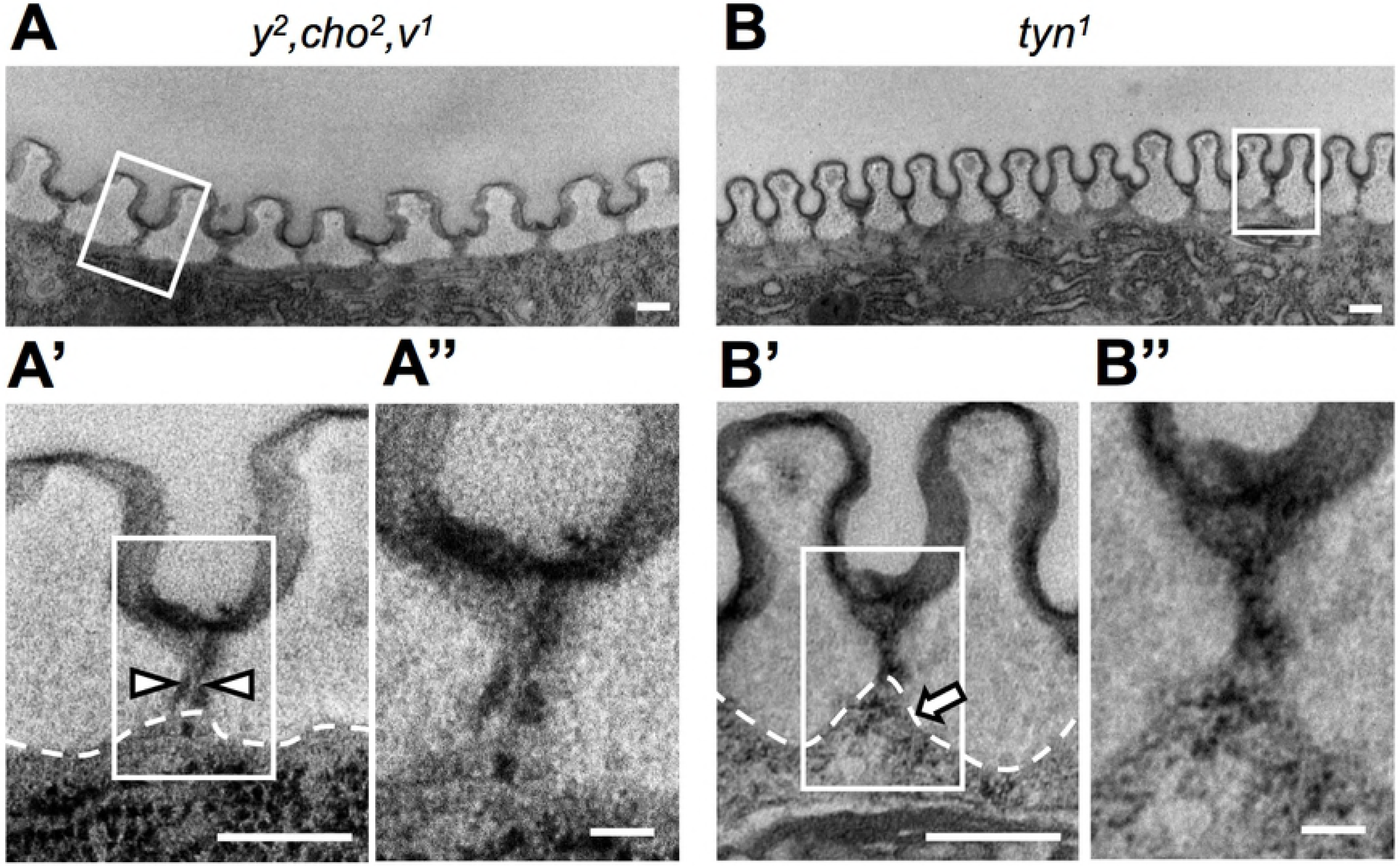
Defective ultrastructures in the *tyn*^*1*^mutant. (A, B) TEM images of the taenidial folds in *y*^*2*^ *, cho*^*2*^ *, v*^*1*^(A-A”) and *tyn*^*1*^ *, y*^*2*^ *, cho*^*2*^ *, v*^*1*^(B-B”). A’, A”, B’ B” are enlarged views of the regions indicated by rectangles in A, A’, B, B’, respectively. Arrowheads in A’ indicate a pore-like structure we called the “taenidial channel” (TC). Dotted lines: plasma membrane. Arrow in B’: a prominent protrusion of the tracheal cell. Scale bars: 200 nm in A, A’, B, B’ and 50 nm in A” and B”.

### Tyn supports the epidermal barrier function

The insect cuticle consists of three layers: the lipid-rich envelope, protein-rich epicuticle, and Chitin-rich procuticle. These layers are thought to play distinct roles in the epidermis. For instance, loss of the *Chitin Synthase-1* (*CS-1*) gene, also called *krotz-kopf verkehrt* (*kkv*), causes an abnormal procuticle, which results in compromised epicuticle, detachment between the cuticle and epidermal cells, and a deformed body shape [35]. Thus, the procuticle layer provides mechanical strength to the cuticle structure. The envelope layer has a barrier function. Loss of the ABC transporter *snustorr* (*snu*) causes an abnormal envelope structure and a dramatically reduced barrier function [26]. The envelope consists of alternating electron-dense and electron-lucid sublayers [7]. In *snu* mutants, one of the electron-dense sublayers is lost. *snu* mutant and RNAi-induced knockdown larvae fail to hatch, and when freed from the egg case, they immediately die from dehydration [26]. In contrast, the *tyn* mutants exhibited a more specific structural anomaly in the envelope, fragmentation of the outermost sublayer. The *tyn* mutant larvae were able to hatch and survive for a few days, despite exhibiting severe behavioral and growth defects, suggesting that dehydration was not the direct cause of their lethality. The Eosin Y permeability in the *tyn* mutants was also much lower than that observed in *snu* mutants [26]. Therefore, some aspects of the barrier function including the internal water retention are maintained in the absence of *tyn* and the outermost envelope sublayer. Our observations using *tyn* mutants support the idea that the outermost envelope layer is essential for the barrier function. The phenotypes of *tyn* and *snu* uncovered non-redundant, protective functions of the envelope sublayers, which together form a robust barrier that protects larvae from the external environment.

### Tyn supports protein clearance from the embryonic trachea

The deposition, assembly, and chemical modification of fibrous aECM consisting of ZP proteins and chitin in the lumen of tracheal tubules are essential for proper regulation of the tube diameter and length [17,21,36-38]. Once the tracheal tubules reach their final shape, the luminal aECM is degraded and replaced by gas prior to larval hatching. A massive wave of endocytosis then removes aECM components into the tracheal cells [33]. At stage 16, prior to this endocytosis wave, 150-500-nm thick cuticle layers develop on the apical surface of tracheal cells [34]. How the degraded luminal aECM is then absorbed efficiently by the tracheal cells through the physical barrier of the cuticles has not been understood. Here, using TEM we observed pore-like structures, which we called “taenidial channels” (TCs), in the inter- taenidial fold region of the tracheal cuticle. The internal surface of the TC was electron-dense and continuous with the envelope layer of the tracheal cuticles, suggesting that it was part of the cuticular envelope. The TCs were abnormal in the *tyn* mutants at the time of the endocytosis wave. We hypothesize that the TC is the channel that permits the passage of materials from the luminal space to the plasma membrane for their efficient endocytic uptake, and that the malformation of TCs in the *tyn* mutants resulted in reduced efficiencies in endocytosis and the clearance of luminal materials.

The *tyn* mutations also delayed and/or inhibited the gas filling of the tracheal system. Although the nature of the gases first appearing in the lumen and the mechanism of gas generation are still not understood, one major hypothesis is that the cavitation forms from gas-saturated luminal liquid on the lipid-covered hydrophobic surface of the cuticle [39,40]. Organic substances remaining in the lumen of *tyn* mutant trachea would reduce the saturation level of gases due to a salting-out effect [41]. Inefficient closure of the tracheal tubule due to defective posterior spiracle formation would further delay gas saturation in the tracheal lumen [19]. These two mechanisms could collectively inhibit the tracheal gas generation in *tyn* mutants.

In the epidermis of insects including *Galleria*, *Tenebrio*, *Tribolium,* and *Drosophila* the pore canal (PC) structure runs vertically through the cuticle layers and was proposed to have a role in wax secretion [42-44]. Similar structures are also described in crustaceans [45]. It will be interesting to determine if the PC has any similarity to the tracheal TC described in this work. The evolutionarily conserved *tyn* gene would be a good starting point for further investigations into the structural and functional characterization of the PC and TC.

## Author Contributions

Y.I. and S.H. conceived the project and designed the experiments. Y.I. and S.I. obtained the experimental data with help from H.W. Y.I. and S.H. analyzed the data and wrote the manuscript.

## Acknowledgements

We thank the Japan National Institute of Genetics and Bloomington Stock Center for providing fly stocks. We also thank Y. Takahashi and S. Yonemura (RIKEN) for generously providing their expert advice and reagents and for allowing us to use their equipment for EM techniques. We are also grateful to T. Nishimura (RIKEN) and members of the Hayashi laboratory for helpful comments. This work was supported in part by “Initiative for the implementation of the diversity research environment” from MEXT JAPAN.

---

**Supplementary Figure 1.** Abnormal larval behavior in response to food supply. Yeast paste was placed at the center of the well at time 0. (Upper) The numbers of *y*^*2*^ *, cho*^*2*^ *, v*^*1*^larvae inside and outside the paste increased and decreased, respectively, meaning that they gradually moved into the yeast paste. Approximately 20% of the larvae were peripheral to the yeast paste at any time point. (Lower) *tyn*^*1*^larvae outside the paste decreased similarly to *y*^*2*^ *, cho*^*2*^ *, v*^*1*^, indicating that the *tyn*^*1*^larvae could sense and move toward the food. However, more larvae tended to stay at the periphery of the food rather than entering it, compared to the control.

**Supplementary Figure 2.** Loss of posterior spiracle valve structures. Posterior spiracles with enlarged views of the posterior tip (dotted-lined region) in the lower right corner are shown for various genotypes (*y*^*2*^ *, cho*^*2*^ *, v*^*1*^, Oregon R, *tyn*^*1*^, *tyn*^*2*^and their sibling controls). The yellow arrows indicate valve structures, which were missing in the *tyn*^*1*^and *tyn*^*2*^mutants.

